# SMAD3 Determines Conventional versus Plasmacytoid Dendritic Cell Fates

**DOI:** 10.1101/715060

**Authors:** Jeong-Hwan Yoon, Eunjin Bae, Katsuko Sudo, Jin Soo Han, Seok Hee Park, Susumu Nakae, Tadashi Yamashita, In-Kyu Lee, Ji Hyeon Ju, Isao Matsumoto, Takayuki Sumida, Masahiko Kuroda, Keiji Miyazawa, Mitsuyasu Kato, Mizuko Mamura

## Abstract

Transforming growth factor (TGF)-β plays crucial roles in differentiation of dendritic cells (DC). However, molecular mechanisms how TGF-β regulates DC differentiation remain largely unknown. Here, we show that selective repression of one of the TGF-β receptor-regulated SMADs (R-SMADs), SMAD3 directs conventional DC (cDC) differentiation, whereas maintenance of SMAD3 is indispensable for plasmacytoid DC (pDC) differentiation. Expression of SMAD3 was specifically downregulated in CD115^+^ common DC progenitor (CDP), pre-cDCs and cDCs. SMAD3 deficient mice showed a significant reduction in pre-pDCs and pDCs with increased CDP, pre-cDCs and cDCs. SMAD3 upregulated the pDC-related genes: SPI-B, E2-2 and IKAROS, while it repressed FLT3 and the cDC-related genes: IRF4 and ID2. STAT3 and a SMAD transcriptional co-repressor, c-SKI repressed SMAD3 for cDC differentiation, whereas canonical SMAD-mediated TGF-β signalling maintained SMAD3 for pDC differentiation. Thus, SMAD3 is the pivotal determinant to bifurcate cDC and pDC differentiation in the steady-state condition.

## INTRODUCTION

A web of cytokine signalling pathways and transcription programs control development of DC subsets from distinct hematopoietic lineages (Lin) (Belz *et al*, 2012; Dress RJ *et al*, 2018; Merad *et al*, 2013; Miller *et al*, 2012; Murphy *et al*, 2016). Several cytokine receptors such as Fms-related tyrosine kinase 3 (FLT3; CD135), c-KIT (CD117) and macrophage colony-stimulating factor receptor (M-CSFR; CD115) are the markers to distinguish Lin^−^ DC progenitors in mouse bone marrow (BM). Common myeloid progenitors (CMPs), common lymphoid progenitor (CLP) and lymphoid-primed multipotent progenitor (LMPP) are the early DC progenitors, which differentiate into the intermediate progenitor, macrophage DC progenitor (MDP). Downstream of MDP is the common DC progenitors (CDPs) comprised of CD115^+^ and CD115^−^ CDPs, which give rise to conventional/classical DC (cDCs) and plasmacytoid DCs (pDCs), respectively, in the steady state condition (Onai *et al*, 2007; Onai *et al*, 2016; Schraml *et al*, 2015). CDPs give rise to Lin^−^CD11c^+^MHCII^−^CD135^+^CD172α^−^ pre-DCs divided into four subsets based on the expression of sialic acid binding Ig-like lectin (Siglec)-H and Ly6C, of which SiglecH^+^ subsets have a pDC potential, while SiglecH^−^ subsets have a cDC potential (Schlitzer *et al*, 2015). Pre-cDCs have been defined as CD11c^+^MHCII^−^ proliferative precursor in BM and lymphoid tissues, which differentiate into CD8^+^ and CD11b^+^ cDCs but not into pDCs (Diao *et al*, 2006; Naik *et al*, 2006). Transcriptome analyses of DC subsets have revealed the essential roles of transcription factors for cDC and pDC differentiation (Miller *et al*, 2012; Murphy *et al*, 2016). The helix-loop-helix (HLH) transcription factor, inhibitor of DNA binding protein 2 (ID2) is required for the development of splenic CD8α^+^ DC subset and Langerhans cells (Hacker *et al*, 2003). Interferon regulatory factors (IRF)-2, IRF-4 and IRF-8 regulate cDC and pDC differentiation (Merad *et al*, 2013). Basic HLH transcription factor (E protein) E2-2/*Tcf4* is a specific transcriptional regulator of pDC, which directly activates pDC-related genes such as *Irf8* and *Spib* (Cisse *et al*, 2008). ID2 and E2-2 induce cDC and pDC, respectively with mutual antagonism (Ghosh *et al*, 2010). The ETS transcription factor, Spi-B expressed in pDC precursors is required for pDC development (Schotte *et al*, 2003, Schotte *et al*, 2004). E2-2 and Spi-B mutually enhance their expressions and cooperate to induce the development of human pDC (Nagasawa *et al*, 2008). IKAROS, a zinc finger protein is required for pDC differentiation by repressing non-pDC gene expression (Allman *et al*, 2006). STAT3 is required for FLT3-dependent DC differentiation (Laouar *et al*, 2003). Although TGF-β has been reported to regulate some of these factors for DC development as introduced below, network of TGF-β signalling pathways and these transcription factors still remains largely to be determined.

TGF-β pleiotropically regulates hematopoiesis and immune cell development (Blank *et al*, 2015; Challen *et al*, 2010; Sanjabi *et al*, 2017; Söderberg *et al*, 2009). TGF-β exerts the differential effects on DC development in ontogenetic stage-dependent manners (Seeger *et al*, 2015). TGF-β promotes DC development from CD34^+^ hemopoietic progenitors (Riedl *et al*, 1997; Strobl *et al*, 1996). TGF-β1 is required for immature DC development, whereas it blocks DC maturation (Yamaguchi *et al*, 1997). TGF-β1 directs differentiation of CDP into cDCs by inducing cDC instructive factors, IRF4 and RelB, and pDC inhibitory factor, ID2 (Felker *et al*, 2010). TGF-β1 induces DC-associated genes such as *Flt3, Irf4* and *Irf8* in multipotent progenitors at steady state (Sere *et al*, 2012). Canonical TGF-β signalling pathway is initiated by ligand-bound activated TGF-β type I receptor (TβRI)-phosphorylated TGF-β receptor-regulated SMADs (R-SMADs): SMAD2 and SMAD3. Despite their high homology, SMAD2 and SMAD3 exert the distinct functions depending on the context (Brown *et al*, 2007; Heldin *et al*, 2012; Massague *et al*, 2005). Signalling mechanisms underlying pleiotropic functions of TGF-β in differentiation of DC subsets from their specific precursors remain unknown.

Here, we have found that SMAD3 is specifically repressed in CD115^+^ CDP, pre-cDCs and cDCs, despite its ubiquitous and constant expression in normal cells in the steady state condition. SMAD3 represses the transcription factors crucial for cDC differentiation such as FLT3, IRF4 and ID2. We have discovered that STAT3 represses SMAD3 for cDC differentiation in cooperation with c-SKI, one of the SKI/SNO protooncoproteins that inhibit TGF-β signalling as the transcriptional corepressor of the SMAD proteins (Deheuninck *et al*, 2009; Massague *et al*, 2005). By contrast, SMAD3 self-induced by SMAD2/3/4 upregulates the pDC-related genes such as Spi-B, E2-2 and IKAROS for pDC differentiation from pre-pDCs, which are substantially deficient in Smad3 null mice. These findings indicate that downregulation or maintenance of SMAD3 expression determines the fate of DC precursors into cDCs and pDCs.

## RESULTS

### Selective downregulation of SMAD3 in cDCs

We examined the basal expression patterns of SMAD2 and SMAD3 in LMPP as Lin^−^Sca-1^+^CD34^+^CD117^+^CD135^+^ cells, DC progenitor cells: MDP as Lin^−^ CD117^hi^CD135^+^CD115^+^Sca-1^−^ cells, CD115^+^ CDP as Lin^−^CD117^int^CD135^+^CD115^+^CD127^−^ cells, CD115^−^ CDP as Lin^−^CD117^int^CD135^+^CD115^−^CD127^−^ cells (Fogg *et al*, 2006; Onai *et al*, 2016), SiglecH^−^Ly6C^−^/SiglecH^−^Ly6C^+^/SiglecH^+^ CD11c^+^MHCII^−^CD135^+^CD172α^−^ pre-DCs (Schlitzer *et al*, 2015) and differentiated DCs: BM CD11c^hi^ cDCs, BM PDCA-1^+^ pDCs, splenic CD11c^hi^ cDCs, splenic PDCA-1^+^ pDCs, lamina propria and Peyer’s patch cDCs of C57BL/6 mice. We also examined GM-CSF plus IL-4-induced BM-derived DCs (BMDCs) that yield CD11b^+^ cDCs and FLT3L-induced BMDCs that yield both cDCs and pDCs (Waskow *et al*, 2008; Yamaguchi *et al*, 1997). SMAD2 mRNA was expressed in all examined cells, whereas SMAD3 mRNA expressed in MDP, CD115^−^ CDP, SiglecH^+^ pre-pDCs and pDCs was reduced to almost undetectable level in CD115^+^ CDP, SiglecH^−^ pre-cDCs and cDCs (Figure 1A and S1A). Immunoblotting confirmed that SMAD2 protein (60 kDa) kept expressed, whereas SMAD3 protein (50 kDa) expressed in whole BM was reduced to undetectable level in GM-CSF plus IL-4-induced BMDCs (Figure S1B).

**Figure 1.**
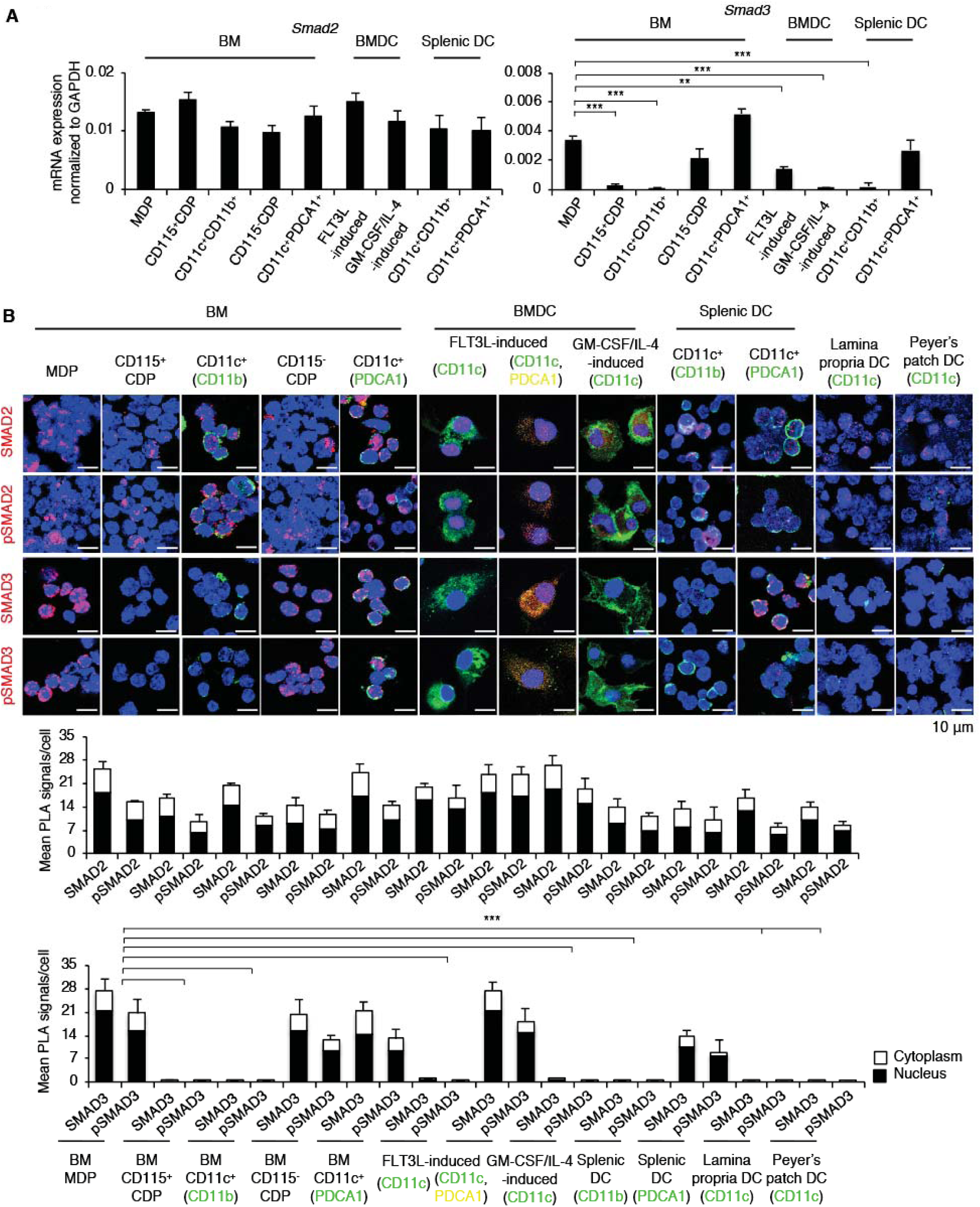
Selective downregulation of SMAD3 in CD115^+^ CDP, pre-cDCs and cDCs. Lin^−^CD117^hi^CD135^+^CD115^+^Sca-1^−^ (MDP), Lin^−^CD117^int^CD135^+^CD115^+^CD127^−^ (CD115^+^ CDP), Lin^−^CD117^int^CD135^+^CD115^−^CD127^−^ (CD115^−^ CDP), SiglecH^−^Ly6C^−^/SiglecH^−^ Ly6C^+^/SiglecH^+^ CD11c^+^MHCII^−^CD135^+^CD172α^−^ (pre-cDC1/ pre-cDC2/ pre-pDC), bone marrow (BM) CD11c^hi^CD11b^+^ cDCs, BM CD11c^+^PDCA-1^+^ pDCs, FLT3L-induced BMDCs, GM-CSF plus IL-4-induced BMDCs, splenic CD11c^hi^CD11b^+^ cDCs, splenic PDCA-1^+^ pDCs were sorted using MACS system and/or FACSAria III. (A) Expression of SMAD2 and SMAD3 mRNA was measured by quantitative RT-PCR. (B) Expression of SMAD2, C-terminally phosphorylated (p) SMAD2, SMAD3 and C-terminally pSMAD3 was measured by proximity ligation assay (PLA). Nucleus was stained with DAPI. CD11c, CD11b and PDCA-1 were stained with Alexa Fluor 488 (green) or 633 (yellow). Red dots in the nucleus (black) and cytoplasm (white) in 10 fields were quantified. Scale bars represent 10 μm. Data are representative of five independent experiments. Graphs show means + s.d. *P* values were calculated by 2-tailed unpaired Student’s *t* test. ** *P*<0.01 and *** *P*<0.0005.

Immunocytochemistry using proximity ligation assay (PLA) showed that SMAD3 protein expressed in LMPP, MDP, CD115^−^ CDP, SiglecH^+^ pre-pDCs and pDCs was not detected in CD115^+^ CDP, SiglecH^−^ pre-cDCs and cDCs, whereas SMAD2 protein was expressed in all subsets (Figure 1B and S1C). SMADs are usually ubiquitously expressed in normal cells (Brown *et al*, 2007) as we observed their expression in naïve and activated CD4^+^ T cells and macrophages (Figure S1D). Thus, SMAD3 is selectively and specifically downregulated in cDCs from their upstream developmental stage at CD115^+^ CDP.

### SMAD3-mediated TGF-β signalling induces pDC while inhibits cDC differentiation

Mutually exclusive expression patterns of SMAD3 in cDCs and pDCs led us to examine the effects of forced expression or knockdown of SMAD3 on DC differentiation using FLT3L-induced or GM-CSF plus IL-4-induced BMDCs transfected with either SMAD3 DNA or SMAD3 siRNA four hours prior to culture. Overexpression of SMAD3 resulted in significant increase in MDP and CD115^−^ CDP while substantial loss of CD115^+^ CDP in FLT3L-induced BMDCs (Figure 2A, upper panels). By contrast, knockdown of SMAD3 significantly decreased MDP, while increased CD115^+^ CDP in both FLT3L-induced or GM-CSF plus IL-4-induced BMDCs (Figure 2A and S2A, lower panels). Forced expression of SMAD3 resulted in significant decrease in MHCII^+^CD11c^+^, CD11b^+^CD11c^+^ cDCs and CD24^+^CD11c^+^ cDCs with significant increase in PDCA-1^+^B220^+^ pDCs in FLT3L-induced BMDCs, whereas knockdown of SMAD3 resulted in significant increase in MHCII^+^CD11c^+^, CD11b^+^CD11c^+^ cDCs and CD24^+^CD11c^+^ cDCs with significant decrease in PDCA-1^+^B220^+^ pDCs in both FLT3L-induced and GM-CSF plus IL-4-induced BMDCs (Figure 2B and S2B). Forced expression of SMAD3 slightly induced pDCs even in GM-CSF plus IL-4-induced BMDCs, which do not develop pDC (Figure S2A and S2B). Forced expression of SMAD3 showed the plasmacytoid morphology with no dendrite formation, whereas knockdown of SMAD3 developed more dendrites compared with control pcDNA or siRNA in BMDCs (Figure 2C and S2C). Knockdown or forced expression of SMAD3 was confirmed by RT-PCR (Figure S2D). SMAD2 and SMAD3 are the R-SMADs shared by TGF-β and Activin among TGF-β superfamily cytokines (Massague *et al*, 2005). Activin A (10 ng/ml) showed no effect on FLT3L or GM-CSF plus IL-4-induced BMDC differentiation (Figure 2D and S2E). High concentration of TGF-β1 (5 ng/ml) completely blocked FLT3L or GM-CSF plus IL-4-induced *Smad3*^+*/*+^ BMDC differentiation, which was abolished in *Smad3*^−*/*−^ BMDCs (Figure 2D and S2E), indicating that potent inhibitory effect of high-dose TGF-β on DC differentiation is SMAD3-dependent. These data show that SMAD3-mediated TGF-β signalling induces pDC differentiation while inhibits cDC differentiation.

**Figure 2.**
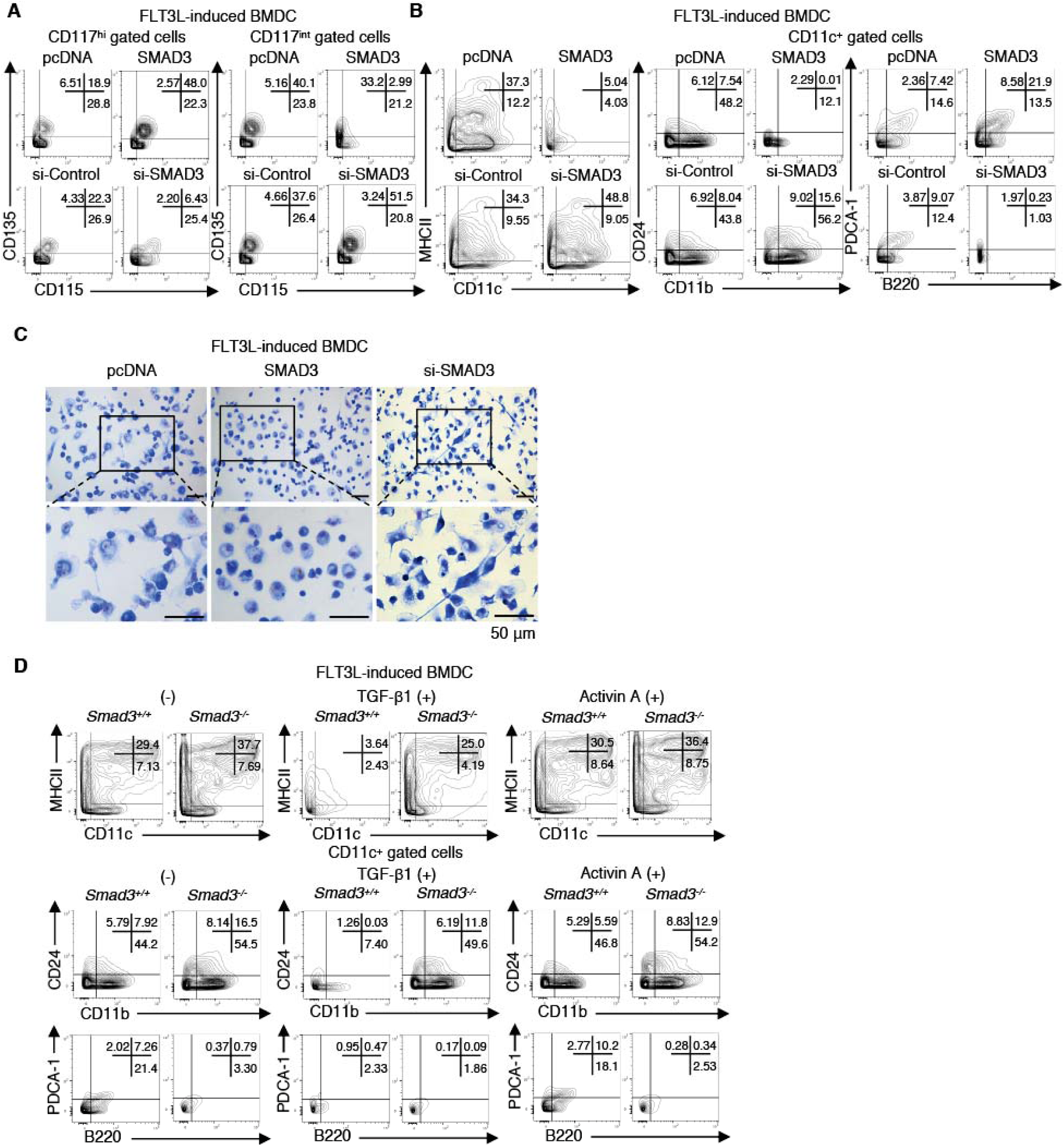
SMAD3 induces pDC, while it inhibits cDC differentiation. FLT3L-induced BMDCs were transfected with SMAD3 DNA, control pcDNA or SMAD3 siRNA, control siRNA 4 hours prior to culture and analyzed on day 7. (A) Contour plots show the expression of CD115 and CD135 on Lin^−^Sca-1^−^CD117^hi^ or Lin^−^ Sca-1^−^CD117^int^ cells. (B) Contour plots show the expression of CD11c/MHCII, CD11b/CD24 and B220/PDCA-1 in CD11c^+^ gate. (C) May-Grunwald/Giemsa stained FLT3L-induced BMDCs transfected with SMAD3 DNA, SMAD3 siRNA or control pcDNA. Scale bars represent 50 μm. (D) Contour plots show the expression of CD11c/MHCII, CD11b/CD24 and B220/PDCA-1 in CD11c^+^ gate of *Smad3*^+*/*+^ or *Smad3*^−*/*−^ FLT3L-induced BMDCs treated with or without TGF-β1 (5 ng/ml) or Activin A (10 ng/ml). Data are representative of three independent experiments.

### Decreased pDCs and increased cDCs in *Smad3*^−*/*−^ mice

We next examined the in vivo effect of SMAD3 deficiency on DC differentiation using *Smad3*^−*/*−^ mice. We confirmed that the proportions of the hematopoietic progenitor cells detected as c-kit^+^Lin^−^Sca-1^+^ (KLS) or CD34^+^ KLS cells, LMPP, CMP as Lin^−^Sca-1^−^CD16/32^−^ CD34^+^CD117^+^ and granulocyte macrophage progenitor (GMP) as Lin^−^Sca-1^−^ CD16/32^+^CD34^+^CD117^+^ were unaltered in BM of 8-weeks old *Smad3*^−*/*−^ mice compared with BM of *Smad3*^+*/*+^ mice bred in specific-pathogen free environment before the onset of any signs of inflammation (Yang *et al*, 1999; Yoon *et al*, 2015) (Figure S3). We observed the significantly increased CD115^+^ CDP with decreased MDP and unaltered CD115^−^ CDP in BM of *Smad3*^−*/*−^ mice compared with BM of *Smad3*^+*/*+^ mice (Figure 3A). DNGR-1 (encoded by the *Clec9a* gene and also known as CLEC9A and CD370) positive CDPs are cDC-restricted (Schraml *et al*, 2013). CX3CR1 is expressed on MDP and cDC-P (Liu *et al*, 2009). *Cx3cr1* and *Dngr1* mRNA expression in Lin^−^CD115^+^ BM cells and CX3CR1^+^CD370^+^Lin^−^ CD117^int^CD135^+^CD115^+^ BM cells were significantly increased in *Smad3*^−*/*−^ mice compared with *Smad3*^+*/*+^ mice (Figure 3B). SiglecH^+^ pre-DCs containing a pDC potential were profoundly reduced, while SiglecH^−^Ly6C^+^ and SiglecH^−^Ly6C^−^ pre-DCs with a cDC potential were significantly increased in BM of *Smad3*^−*/*−^ mice compared with BM of *Smad3*^+*/*+^ mice (Figure 3C). Total cDCs in BM, spleen, superficial and mesenteric lymph nodes of *Smad3*^−*/*−^ mice were significantly increased compared with those of *Smad3*^+*/*+^ mice, whereas SiglecH^+^PDCA1^+^B220^+^ pDCs were substantially undetected in those of *Smad3*^−*/*−^ mice (Figure 3D). Immunophenotyping of DC subsets in *Smad3* deficient mice shows that SMAD3 deficiency facilitates cDC differentiation at the developmental stage between MDP and CD115^+^ CDP, while it blocks differentiation of pre-pDCs and pDCs in the steady state.

**Figure 3.**
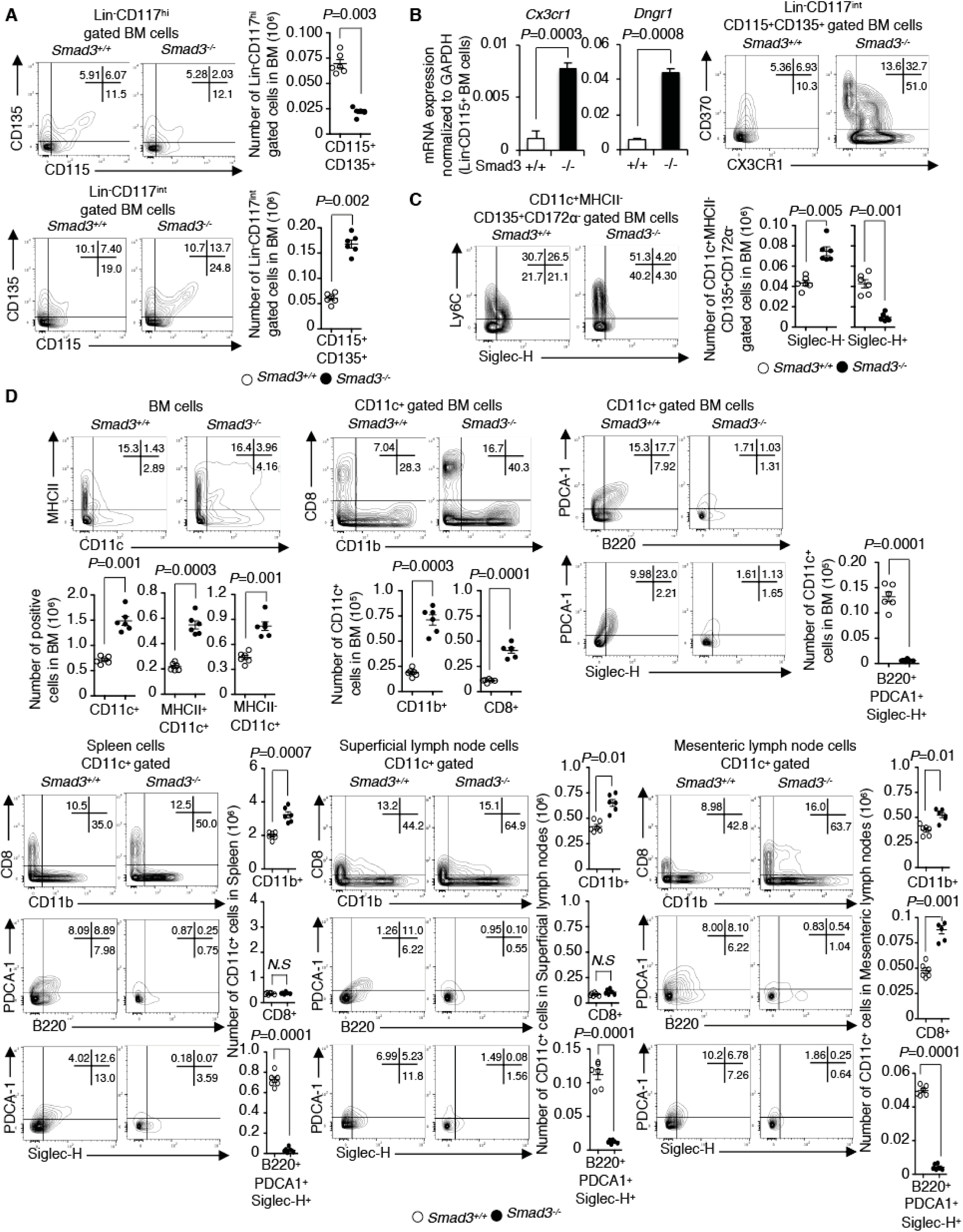
Lack of pDC with increase of cDC in *Smad3*^−*/*−^ mice. Immunophenotyping of *Smad3*^−*/*−^ or *Smad3*^+*/*+^ mice (n = 6/genotype) was performed using flowcytometry. (A) Contour plots show the expression of CD115/CD135 in Lin^−^Sca-1^−^CD117^hi^ or Lin^−^Sca-1^−^ CD117^int^ gates. Graphs show the cell numbers of Lin^−^CD117^hi^CD115^+^CD135^+^ MDP and Lin^−^ CD117^int^CD115^+^CD135^+^ CDP in BM. (B) Quantitative RT-PCR of *Cx3cr1* and *Dngr1* mRNA in Lin^−^CD115^+^ BM cells and contour plots of CX3CR1^+^CD370^+^Lin^−^CD117^int^CD115^+^CD135^+^ BM cells. (C) Contour plots show the expression of SiglecH/Ly6C in CD11c^+^MHCII^−^CD135^+^CD172α^−^ pre-DC gate. Graphs show the cell numbers in BM. (D) Contour plots show the expression of CD11c, MHCII, CD11b, CD8, SiglecH, PDCA-1 and B220 in BM, spleen, superficial and mesenteric lymph nodes. Graphs show the cell numbers of CD11b^+^, CD8^+^, SiglecH^+^PDCA-1^+^B220^+^ cells in CD11c^+^ gate. Graphs show means ± s.d. *P* values were calculated by 2-tailed unpaired Student’s *t* test.

### SMAD3 represses cDC-related genes while induces pDC-related genes

To identify the responsible genes to facilitate cDC while to inhibit pDC differentiation in *Smad3*^−*/*−^ mice, we compared the mRNA expressions of the cytokines, their signalling molecules and the transcription factors essential for DC differentiation in DC precursors and subsets of *Smad3*^−*/*−^ mice with those of littermate control *Smad3*^+*/*+^ mice. It was impossible to sort out pre-pDCs and pDCs from BM and lymphoid organs of *Smad3*^−*/*−^ mice because they were substantially deficient (Figure 3C and 3D). *Smad3*^−*/*−^ CD115^+^ CDPs, pre-cDCs and cDCs expressed significantly higher levels of cDC-related genes: *Flt3, Irf4* and *Id2* compared with *Smad3*^+*/*+^ controls, whereas *Smad3*^−*/*−^ and *Smad3*^+*/*+^ CD115^−^ CDPs expressed similar levels of pDC markers: *Siglech* (Takagi *et al*, 2011), *Bst2* encoding PDCA-1 (Blasius *et al*, 2006) and the transcription factors essential for pDC differentiation: *Spib, E2-2* and *Ikaros* (Figure 4A). To identify the target genes for SMAD3 to induce pDC while to inhibit cDC differentiation, we screened these genes in BMDCs transfected with either SMAD3 DNA or control pcDNA. We found that forced expression of SMAD3 significantly upregulated the mRNA expression of pDC-related genes: *Spib, E2-2* and *Ikaros* in FLT3L-induced BMDCs (Figure 4B and S4A). SMAD3 slightly upregulated these pDC-related genes even in GM-CSF plus IL-4-induced BMDCs (Figure S4A and S4B). Low concentrations of TGF-β1 (0.1, 0.2 ng/ml) showed the same effect with forced expression of SMAD3 on these pDC-related genes in both FLT3L-induced and GM-CSF plus IL-4-induced *Smad3*^+*/*+^ BMDCs, which was abolished in *Smad3*^−*/*−^ BMDCs (Figure 4C and S4C). By contrast, forced expression of SMAD3 significantly repressed the mRNA expression of cDC-related genes: *Flt3, Irf4* and *Id2* in FLT3L-induced BMDCs, *Flt3* and *Irf4* in GM-CSF plus IL-4-induced BMDCs (Figure 4D, S4A and S4D). Low concentrations of TGF-β1 repressed these cDC-related genes in *Smad3*^+*/*+^ BMDCs, which remained in *Smad3*^−*/*−^ BMDCs (Figure 4E and S4E). High concentrations of TGF-β1 completely repressed both cDC and pDC-related genes in *Smad3*^+*/*+^ BMDCs (Figure 4C and 4E). Thus, SMAD3-mediated low-dose TGF-β signalling induces pDC-related genes while represses cDC-related genes.

**Figure 4.**
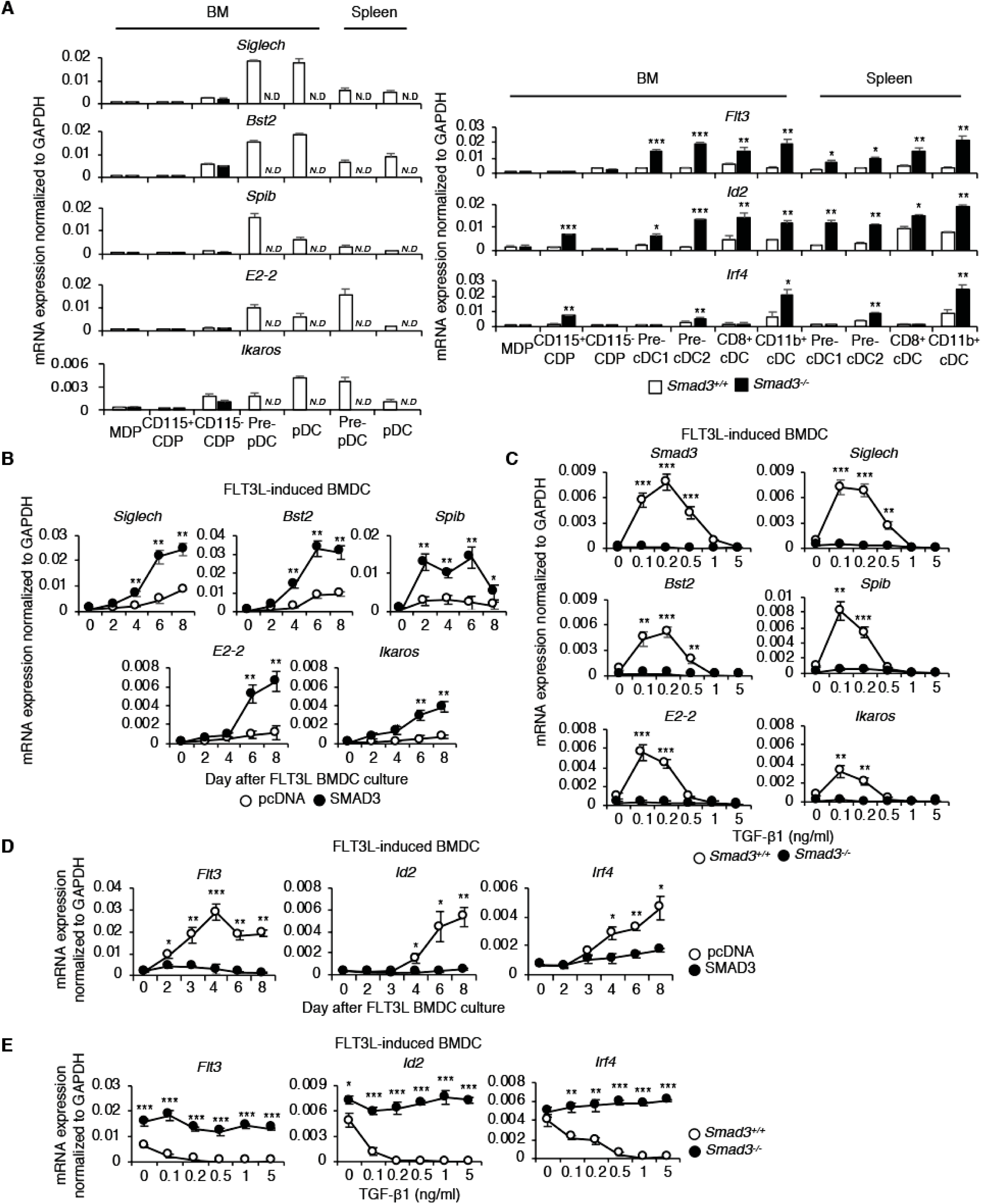
SMAD3 represses cDC-related genes while induces pDC-related genes. Expressions of *Siglech, Bst2, Spib, E2-2, Ikaros, Flt3, Id2* and *Irf4* mRNA in (A) MDP, CD115^+^ CDP, CD115^−^ CDP, pre-cDC1/ pre-cDC2/ pre-pDC, BM CD11c^hi^ cDCs, BM pDCs, FLT3L-induced BMDCs, GM-CSF plus IL-4-induced BMDCs, splenic cDCs, splenic pDCs from *Smad3*^−*/*−^ or *Smad3*^+*/*+^ mice. Expressions of *Siglech, Bst2, Spib, E2-2, Ikaros* mRNA in (B) FLT3L-induced BMDCs transfected with SMAD3 DNA or control pcDNA 4 hours prior to culture and analyzed on day 2, 4, 6, 8 and in (C) *Smad3*^−*/*−^ or *Smad3*^+*/*+^ FLT3L-induced BMDCs treated with the indicated concentrations of TGF-β1 on day 8 were determined by quantitative RT-PCR. Expressions of *Flt3, Id2* and *Irf4* mRNA in (D) FLT3L-induced BMDCs transfected with SMAD3 DNA or control pcDNA 4 hours prior to culture on day 2, 3, 4, 6, 8 and in (E) *Smad3*^−*/*−^ or *Smad3*^+*/*+^ FLT3L-induced BMDCs treated with the indicated concentrations of TGF-β1 on day 8 were determined by quantitative RT-PCR. Data are representative of three independent experiments with triplicate. Graphs show means ± s.d. *P* values were calculated by 2-tailed unpaired Student’s *t* test. * *P*<0.05, ** *P*<0.01 and *** *P*<0.0005.

### TGF-β induces transactivation of the *Smad3* gene, thereby pDC differentiation

Because low concentration of TGF-β upregulated pDC-related genes while repressed cDC-related genes, we examined the dose effect of TGF-β on DC differentiation. In accordance with the effect on the gene expression, low concentrations of TGF-β1 significantly increased pDC differentiation while suppressed cDC differentiation in FLT3L-induced BMDCs, which was abolished in *Smad3*^−*/*−^ BMDCs (Figure 5A). Low concentrations of TGF-β1 induced pDC differentiation even in GM-CSF plus IL-4-induced *Smad3*^+*/*+^ BMDCs, which was abolished in *Smad3*^−*/*−^ BMDCs (Figure 5B). High concentrations of TGF-β1 potently inhibited both pDC and cDC differentiation (Figure 5A and 5B). SMAD3 was C-terminally phosphorylated in the nuclei in CD115^−^ CDP, pre-pDCs and pDCs (Figure 1B and S1C). PLA showed that low concentrations of TGF-β1 upregulated the expression and C-terminal phosphorylation levels of SMAD3 proteins (Figure 5C). These data verify that SMAD3-mediated low-dose TGF-β signalling induces pDC while represses cDC differentiation.

**Figure 5.**
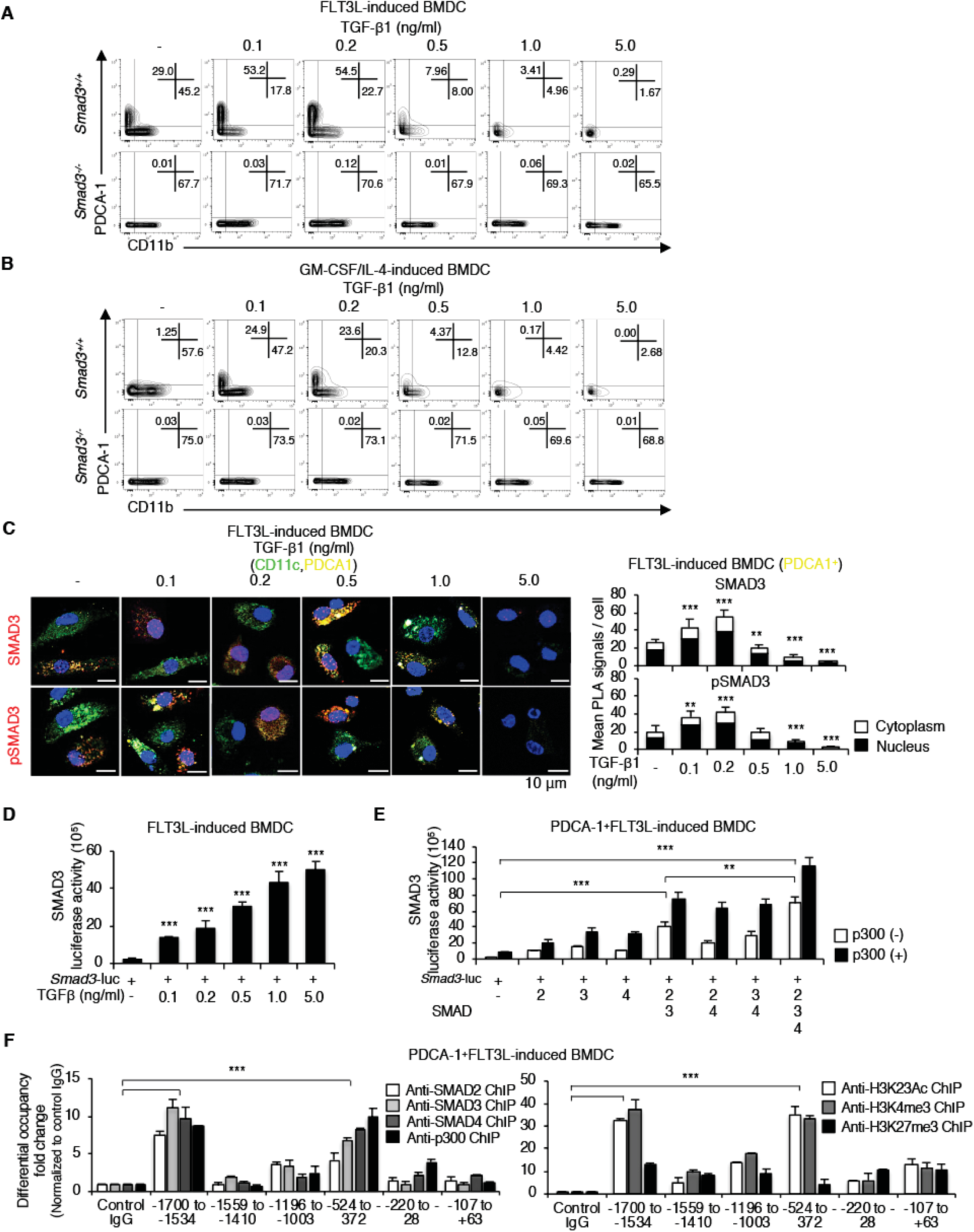
TGF-β induces transcription of the *Smad3* gene, thereby pDC differentiation. Contour plots show the expressions of CD11b/PDCA-1 in CD11c^+^ gate of (A) FLT3L-induced BMDCs and (B) GM-CSF plus IL-4-induced BMDCs from *Smad3*^−*/*−^ or *Smad3*^+*/*+^ mice treated with the indicated concentrations of TGF-β1. Data are representative of three independent experiments. (C) Expressions of SMAD3 and C-terminally pSMAD3 in FLT3L-induced BMDCs treated with the indicated concentrations of TGF-β1 were determined by PLA (red). Nucleus was stained with DAPI. CD11c was stained with Alexa Fluor 488. PDCA-1 was stained with Alexa Fluor 633. Red dots in the nucleus (black) and cytoplasm (white) in 10 fields were quantified. Scale bars represent 10 μm. (D) Smad3 promoter activity was determined using FLT3L-induced BMDCs transfected with the *Smad3* promoter luciferase reporter construct treated with the indicated concentrations of TGF-β1. (E) Smad3 promoter activity was determined using PDCA-1^+^FLT3L-induced BMDCs transfected with the *Smad3* promoter luciferase reporter construct with the indicated combinations of SMAD2, SMAD3, SMAD4 and p300. (F) Binding of SMAD2, SMAD3, SMAD4, p300 and the histone modification in the *Smad3* proximal promoter region in PDCA-1^+^FLT3L-induced BMDCs was determined by chromatin immunoprecipitation (ChIP) using the antibodies against SMAD2, SMAD3, SMAD4, p300, acetyl histone H3K23 (H3K23Ac), trimethyl histone H3K4 (H3K4me3) and trimethyl histone H3K27 (H3K27me3). FLT3L-induced BMDCs were transfected with the *Smad3* promoter luciferase reporter construct and the indicated DNA constructs 4 hours prior to culture and analyzed on day 7. Data of the *Smad3* promoter luciferase reporter assay and ChIP are representative of three to five independent experiments with triplicate. Graphs show means + s.d. *P* values were calculated by 2-tailed unpaired Student’s *t* test. ** *P*<0.01 and *** *P*<0.0005.

To investigate the mechanisms how TGF-β signalling maintains the *Smad3* mRNA expression for pDC differentiation, we examined the effect of SMAD-mediated TGF-β signalling on the *Smad3* gene promoter activity using the luciferase reporter spanning 2 kilobase upstream of the first exons of the *Smad3* genes transfected in FLT3L-induced BMDCs. TGF-β1 induced the *Smad3* promoter activity in a dose-dependent manner (Figure 5D). Discrepancy between lower SMAD3 protein expression and the dose dependency observed in luciferase assay in FLT3L-induced BMDCs treated with the high concentrations of TGF-β1 (0.5, 1.0, 5.0 ng/ml) might be due to the potent inhibition of total DC differentiation (Figure 2D and S2E) and the ligand-dependent degradation of SMAD3 (Fukuchi *et al*, 2001). SMAD2, SMAD3, SMAD4 and their transcription coactivator and histone acetyl-transferase p300 (Massague *et al*, 2005; Heldin *et al*, 2012) synergistically induced the *Smad3* promoter activity as well as TGF-β1 (Figure 5E). Chromatin immunoprecipitation (ChIP) showed that SMAD2, SMAD3, SMAD4 and p300 were bound to the same sites in PDCA1^+^ cells isolated from FLT3L-induced BMDCs (−1700 to −1534 and −524 to −372), which were epigenetically active with acetyl histone H3K23 and trimethyl histone H3K27 (Figure 5F). Taken together, transactivation of SMAD3 via canonical TGF-β signalling pathway initiated by low-dose TGF-β induces the essential transcription factors for pDC differentiation.

### STAT3 and c-SKI repress transcription of SMAD3 for cDC differentiation

We next investigated the mechanisms how SMAD3 is downregulated in CD115^+^ CDP for cDC differentiation. The ligand of CD115, M-CSF induces STAT3 activation in macrophages (Novak *et al*, 1995). STAT3 functions as the signalling molecule of FLT3L and GM-CSF (Li *et al*, 2013; Onai *et al*, 2006; Wan *et al*, 2013) and is essential for FLT3L-responsive DC progenitor proliferation (Laouar *et al*, 2003). Thus, we examined the effect of STAT3 on SMAD3 expression in BMDCs transfected with STAT3 and control pcDNA or siSTAT3 and control siRNA. Forced expression of STAT3 suppressed, whereas knockdown of STAT3 upregulated the expression of *Smad3* mRNA (Figure 6A and S5A). We examined whether and how STAT3 regulates the *Smad3* gene promoter activity using the *Smad3* gene promoter luciferase reporter construct transfected in FLT3L-induced or GM-CSF plus IL-4-induced BMDCs. CD11b^+^ cells were sorted as cDCs before cell lysis for FLT3L-induced BMDCs. SMAD2 and SMAD3 synergistically induced the *Smad3* promoter activity, which was repressed by STAT3 (Figure 6B and S5B). We screened the representative transcriptional repressors of R-SMADs: SKI/SnoN and TGIF as a corepressor of STAT3 (Deheuninck *et al*, 2009; Heldin *et al*, 2012; Massague *et al*, 2005). Among them, c-SKI showed the synergy with STAT3 to repress the *Smad3* promoter activity (Figure S5C). Knockdown or forced expression of c-SKI and STAT3 showed the same effect on *Smad3* mRNA expression in FLT3L-induced and GM-CSF plus IL-4-induced BMDCs (Figure 6A and S5A). Knockdown of c-SKI by siRNA completely abolished the repressive effect of STAT3 on the SMAD2/3-induced *Smad3* promoter activation (Figure 6C and S5D). By contrast, c-SKI alone retained the repressive effect on the SMAD2/3-induced *Smad3* promoter activation when STAT3 was knocked-down, although the synergistic repressive effect of c-SKI and STAT3 was more effective (Figure 6D). ChIP showed that SMAD2 was bound to the same sites with STAT3 and c-SKI in FLT3L-induced BMDCs (−1196 to −1003 and −220 to −28), which were epigenetically inactive with trimethyl histone H3K27 (Figure 6E). These data indicate that STAT3 in synergy with c-SKI represses SMAD-induced transcription of the *Smad3* gene for cDC differentiation.

**Figure 6.**
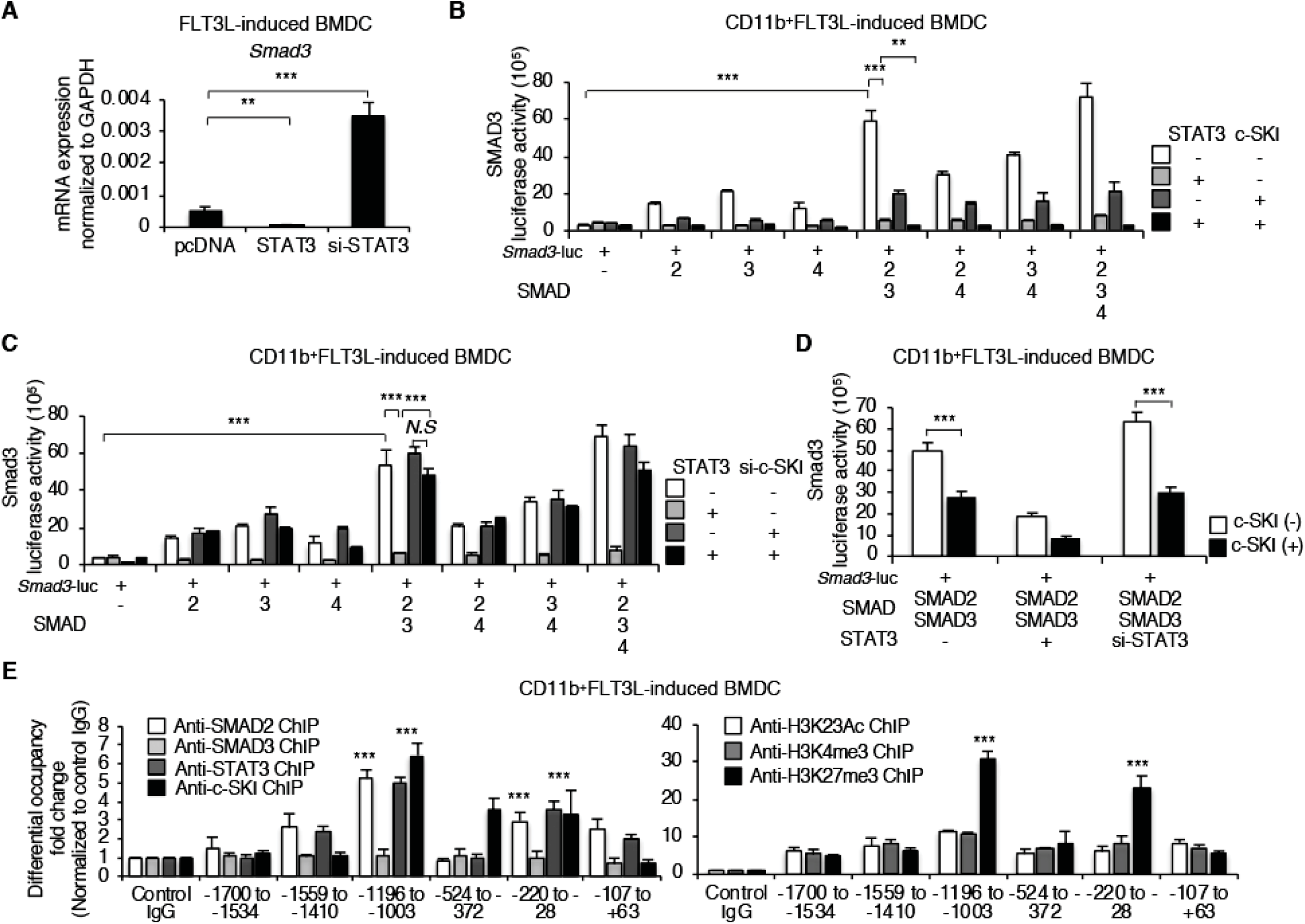
STAT3 and c-SKI repress transcription of SMAD3 for cDC differentiation. (A) Expressions of *Smad3* mRNA in FLT3L-induced BMDCs transfected with STAT3 DNA, control pcDNA or STAT3 siRNA were determined by quantitative RT-PCR. Smad3 promoter activity was determined using CD11b^+^FLT3L-induced BMDCs transfected with the *Smad3* promoter luciferase reporter construct with the indicated combinations of (B) SMAD2, SMAD3, SMAD4, STAT3 and c-SKI, (C) SMAD2, SMAD3, SMAD4, STAT3 and c-SKI siRNA, (D) SMAD2, SMAD3, STAT3 siRNA and c-SKI. (E) Binding of SMAD2, SMAD3, STAT3, c-SKI and the histone modification in the *Smad3* proximal promoter region in CD11b^+^FLT3L-induced BMDCs was determined by ChIP using the antibodies against SMAD2, SMAD3, STAT3, c-SKI, H3K23Ac, H3K4me3 and H3K27me3. FLT3L-induced BMDCs were transfected with the *Smad3* promoter luciferase reporter construct, the indicated siRNA and DNA constructs 4 hours prior to culture and analyzed on day 7. *P* values were calculated by 2-tailed unpaired Student’s *t* test. ** *P*<0.01 and *** *P*<0.0005.

### Interaction of phosphorylated STAT3 with c-SKI is essential for repression of SMAD3 in cDC

We sought to confirm the physiological interactions between STAT3, c-SKI and SMAD2 in DC progenitors, pre-DCs, DC subsets, FLT3L-induced and GM-CSF plus IL-4-induced BMDCs. PLA showed the interaction of c-SKI with STAT3 and SMAD2 in CD115^+^ CDP, pre-cDCs, cDCs and CD11c^hi^ BMDCs, but not in the early progenitors, pre-pDCs and pDCs (Figure 7A and S6A). Knockdown of c-SKI by siRNA abolished the interaction of SMAD2 and STAT3, whereas knockdown of STAT3 had no effect on the interaction of c-SKI and SMAD2 in CD11c^hi^ BMDCs (Figure 7B).

**Figure 7.**
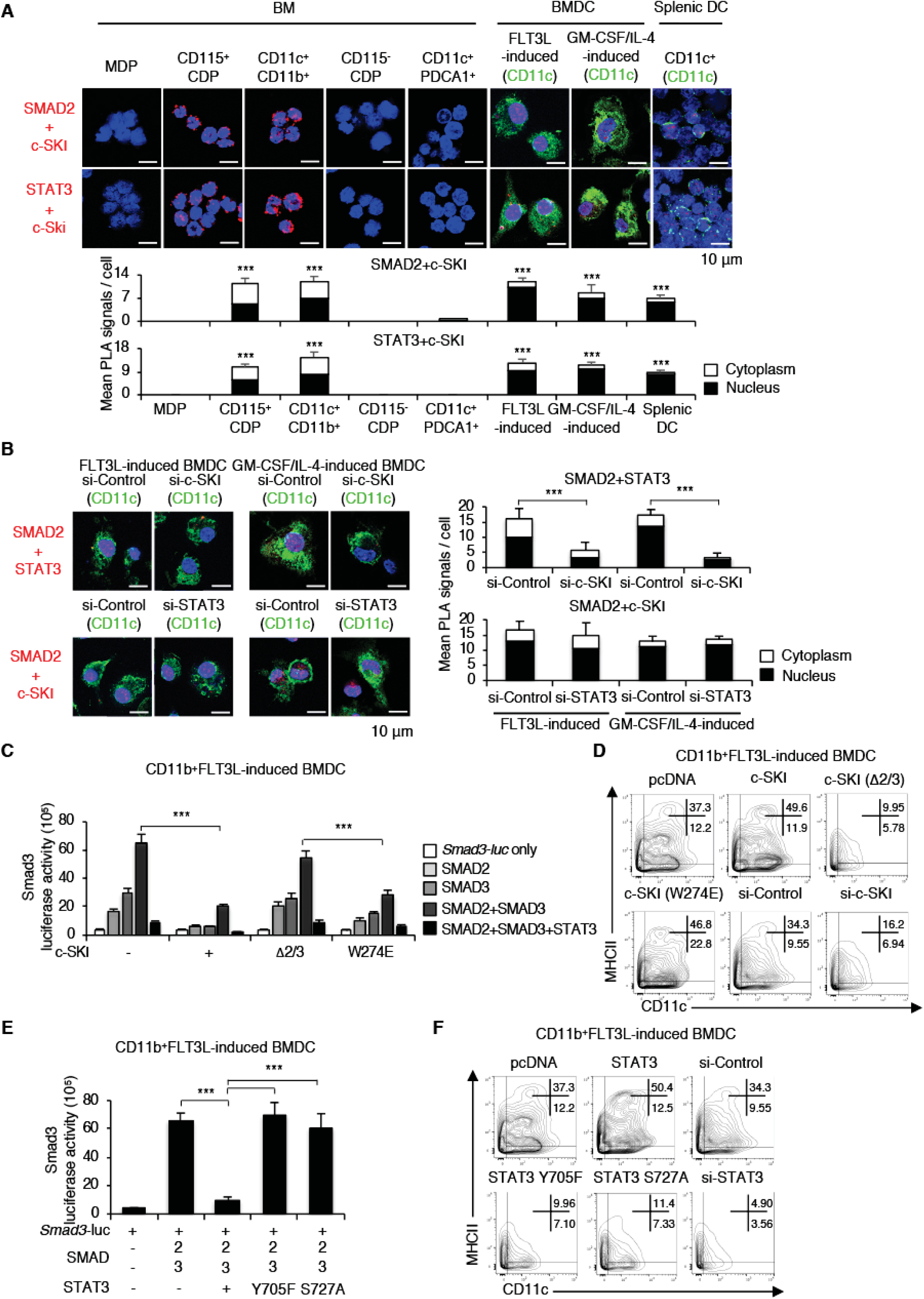
Interaction of phosphorylated STAT3 with c-SKI is essential for repression of SMAD3 in cDC. Proximity between (A) SMAD2 and c-SKI, c-SKI and STAT3 in MDP, CD115^+^ CDP, BM CD11c^hi^CD11b^+^ cDCs, CD115^−^ CDP, BM pDCs, FLT3L-induced BMDCs, GM-CSF plus IL-4-induced BMDCs, splenic cDCs, (B) STAT3 and SMAD2, SMAD2 and c-SKI in FLT3L-induced or GM-CSF plus IL-4-induced BMDCs transfected with the indicated c-SKI siRNA, STAT3 siRNA or control siRNA was determined by PLA. Nucleus was stained with DAPI. CD11c was stained with Alexa Fluor 488 (green). Red dots in the nucleus (black) and cytoplasm (white) in 10 fields were quantified. Scale bars represent 10 μm. Data are representative of three independent experiments. Graphs show means + s.d. (C) Effects of c-SKI mutations (Δ2/3 that does not interact with Smad2/3 and W274E that does not interact with Smad4) on *Smad3* promoter activity in CD11b^+^FLT3L-induced BMDCs were determined by the *Smad3* promoter luciferase reporter assay. (D) Contour plots show the expression of CD11c/MHCII in CD11b^+^gate of FLT3L-induced BMDCs transfected with the indicated c-SKI mutants, c-SKI siRNA or control pcDNA, control siRNA. (E) Effects of STAT3 phosphorylation site-specific mutants (Y705F and S727A) on SMAD2/3-induced activation of the *Smad3* promoter constructs transfected in FLT3L-induced BMDCs were determined by the *Smad3* promoter luciferase reporter assay. (F) Contour plots show the expression of CD11c/MHCII in CD11b^+^gate of FLT3L-induced BMDCs transfected with the indicated STAT3 phosphorylation site-specific mutants (Y705F and S727A) or STAT3 siRNA, control siRNA 4 hours prior to culture and analyzed on day 7. Data are representative of three independent experiments. Graphs show means + s.d. *P* values were calculated by 2-tailed unpaired Student’s *t* test. ** *P*<0.01 and *** *P*<0.0005.

We next investigated the mechanisms how c-SKI and STAT3 repress transcription of the *Smad3* gene using the *Smad3* gene promoter luciferase reporter construct with various combinations of mutants of c-SKI and STAT3 transfected in CD11b^+^FLT3L-induced or GM-CSF plus IL-4-induced BMDCs. SKI interacts with SMAD2 and SMAD3 through its N-terminal region or with SMAD4 through its SAND-like domain to block the ability of the SMAD complexes to activate transcription of TGF-β target genes (Akiyoshi *et al*, 1999; Massague *et al*, 2005; Suzuki *et al*, 2004; Takeda *et al*, 2004). A mutant of c-SKI that does not interact with SMAD2/3 (Δ2/3) failed to repress the *Smad3* promoter activity, whereas a mutant of c-SKI that does not interact with SMAD4 (W274E) (Wu *et al*, 2002) retained the repressive effect on the *Smad3* promoter activity (Figure 7C and S6B). Transfection of Δ2/3 and c-SKI siRNA failed to induce cDC differentiation, whereas W274E significantly upregulated cDC differentiation as well as wild type c-SKI (Figure 7D and S6C). These data verify that interaction of c-SKI with SMAD2 and SMAD3 is required for STAT3-mediated repression of SMAD3 and cDC differentiation. STAT3 is phosphorylated at Y705 and S727 residues upon stimulation (Hillmer *et al*, 2016). Inactive mutants of STAT3 at Y705 and S727 residues, Y705F and S727A, respectively, abolished the repressive effect on SMAD2/3-induced *Smad3* promoter activition in CD11b^+^FLT3L-induced or GM-CSF plus IL-4-induced BMDCs (Figure 7E and S6D). Forced expression of STAT3 signifiantly upregulated cDC differentiation to the same degree as forced expression of c-SKI, which was abolised by Y705F and S727A mutations to the same degree as STAT3 siRNA (Figure 7F and S6E). Thus, c-SKI is necessary for phosphorylated STAT3 at Y705 and S727 to interact with TGF-β R-SMADs to repress transcription of the *Smad3* gene for cDC differentiation.

## DISCUSSION

Major DC subsets are classified into cDCs and pDCs, even though recent studies have shown increasingly complex classification of DC subpopulations and progenitors during ontogeny (Dress *et al*, 2018). Signalling mechanisms how TGF-β regulates the differentiation of DC subsets in the steady state remained largely unknown, although its importance in immunogenic and tolerogenic functions of the differentiated DCs have been appreciated (Seeger *et al*, 2015). In this study, specific repression of SMAD3 in cDCs and their progenitors: pre-cDCs and CD115^+^ CDP led us to uncover its pivotal role to bifurcate DC differentiation into cDCs and pDCs.

SMAD2 and SMAD3 are ubiquitously and constitutively expressed in normal cells in general, whereas their loss is frequently observed in various cancers (Brown *et al.*, 2007; Heldin *et al.*, 2012). They have high amino acid sequence identity in their MH2 domains containing two C-terminal serine residues, 465/467; nevertheless they regulate the same or distinct sets of TGF-β target genes to exert redundant or distinct functions depending on the context (Brown *et al*, 2007, Massague *et al*, 2005). However, the precise mechanisms how they are selected for distinct functions by the context still remain largely unknown. In order to prevent continuous SMAD-mediated TGF-β signalling in normal epithelial cells, SMAD2 and SMAD3 are negatively regulated by diverse mechanisms (Brown *et al*, 2007). TGF-β has been shown to downregulate SMAD3 in lung epithelial cells through decreased transcription (Yanagisawa *et al*, 1998) or in epitheilial-to mesenchymal transition of human glomerular mesangial cells through decreased transcription and increased ubiquitination (Poncelet *et al*, 2007), suggesting that downregulation of SMAD3 can be one of the mechanisms to select the appropriate R-SMAD in response to TGF-β by the context. This study is the first to show that the transcriptional downregulation of SMAD3 is indispensable for differentiation of the normal immune cell subsets in the steady-state. Further studies are required to explore the distinct regulation of R-SMADs in immune cells including the effector DC subsets in the pathophysiological setting (Merad *et al*, 2013).

This study shows that STAT3 transcriptionally represses SMAD3 to derepress cDC-related genes: IRF4, ID2 and FLT3 in cDCs, pre-cDC and CD115^+^ CDP. Indispensable role of STAT3 in DC development has been shown by loss of cDCs by deletion of STAT3 in vivo (Laouar *et al*, 2003) and promotion of DC maturation from the progenitors by overexpression of STAT3 (Onai *et al*, 2006). Engagement of FLT3L and FLT3 expressed on DC precursors such as MDP, CDPs, and pre-cDCs (Liu *et al*, 2009; Onai *et al*, 2007) induces rapid phosphorylation of STAT3 and FLT3L-responsive DC progenitor proliferation (Li *et al*, 2013). STAT3 transiently activated by GM-CSF promotes differentiation of myeloid lineages including cDCs (Merad *et al*, 2013; Wan *et al*, 2013). Colony stimulating factor, the ligand of CD115 induces STAT3 activation (Novak *et al*, 1995). This study has revealed the previously unknown mechanism how STAT3 induces cDC differentiation through repressing SMAD3, a repressor of the representative cDC-related genes.

We have further elucidated that c-SKI is required for STAT3 to repress SMAD3 for cDC differentiation. SKI and the closely related SnoN oncogenes act as transcriptional co-repressors in TGF-β signalling through interaction with SMADs (Akiyoshi *et al*, 1999; Suzuki *et al*, 2004; Wu *et al*, 2002). Although SKI is more widely expressed than SnoN in mature hematopoietic cells and play crucial roles in hematopoiesis and myeloproliferative diseases (Pearson-White *et al*, 1995; Singbrant *et al*, 2014), very little was known about its roles in differentiation and functions of immune cells. We have previously shown that SKI and SnoN oncoproteins cooperate with phosphorylated STAT3 in an adenocarcinoma lung cancer cell line, HCC827 to repress transcription of the *Smad3* gene, which renders the sensitive cells resistant to gefitinib (Makino *et al*, 2017). Here, we have found that c-SKI, but not SnoN is indispensable for STAT3 to repress the *Smad3* gene for cDC differentiation rather than playing a conventional role as a transcriptional co-repressor of SMADs. In hematopoietic cells, SKI represses retinoic acid receptor signalling (Dahl *et al*, 1998), which enhances SMAD3/SMAD4-driven transactivation (Pendaries *et al*, 2003). SKI induces a gene signature associated with hematopoietic stem cells (HSCs) and myeloid differentiation, as well as hepatocyte growth factor (HGF) signalling (Singbrant *et al*, 2014). These previous reports and our finding suggest the possibility that HGF signals via STAT3 (Schaper *et al*, 1997) might induce synergy with c-SKI to repress SMAD3 toward myeloid differentiation. Distinct binding sites of SMADs, p300, STAT3 and c-SKI in the *Smad3* promoter in pDCs and cDCs shown in this study and HCC827 lung cancer cell line are consistent with the previous report showing that cell-type-specific master transcription factors direct SMAD3 to distinct specific binding sites to determine cell-type-specific responses to TGF-β signaling (Mullen et al, 2011).

Studies prior to the identification of distinct CDP subsets (Onai *et al*, 2013) have shown that TGF-β facilitates cDC differentiation from CDP by inducing the essential factors for cDC differentiation such as IRF4, IRF8, RelB, ID2 and FLT3 (Felker *et al*, 2010; Sere *et al*, 2012). They have reported that TGF-β induces ID2 (Hacker *et al*, 2003). By contrast, TGF-β represses ID2 in epithelial cells (Zavadil *et al*, 2005), which is consistent with our data. The discrepancy between these previous reports and this study may be attributed to their two-step amplification and differentiation culture systems. The first-step amplification culture contains SCF, hyper-IL-6 and insulin-like growth factor-1, which modulate STAT and SMAD signalling pathways (Rojas *et al*, 2016; Sarenac *et al*, 2016; Yoon *et al*, 2015). They report that the amplified DC precursor cells with these cytokines prior to DC differentiation are CDP (Felker *et al*, 2010), which may be regulated by TGF-β through a mechanism distinct from the upstream progenitor cells. The second-step differentiation culture contains GM-CSF and IL-4. However, neither TGF-β nor forced expression and knockdown of SMAD3 affected the expression of ID2 in GM-CSF plus IL-4-induced BMDCs in our system. Moreover, the expression of ID2 is not downregulated as deduced from these reports, but rather significantly upregulated in cDCs, pre-cDCs and CD115^+^ CDP in *Smad3*^−*/*−^ mice. These data support the pre-selection of committed CDP subset(s) in the first-step culture containing FLT3L in these previous reports.

In contrast with the repressive effect on cDC-related genes, SMAD3 induces pDC-related genes. Consistently, we have found that SMAD3 deficiency in vivo resulted in substantial loss of pDCs and pre-pDCs, indicating the indispensable role of SMAD3 for pDC differentiation at the developmental stage of pre-pDCs. We have found that low-dose TGF-β in vitro that represents in vivo steady-state physiological function (Zi *et al*, 2012) transactivates SMAD3 in BMDCs, which then induces pDC-related genes: *Spib, Tcf4* and *Ikaros* (Allman *et al*, 2006; Cisse *et al*, 2008; Ghosh *et al*, 2010; Nagasawa *et al*, 2008, Reizis *et al*, 2019). Significantly more profound decrease of pDCs in SMAD3 deficient mice compared with the mice deficient in E2-2 (Ghosh *et al*, 2010; Grajkowska *et al*, 2017) also supports our finding that SMAD3 is the upstream inducer of these pDC-related genes. The proximal promoter regions of these pDC-related genes and pDC marker genes: *Siglech, Bst2* upregulated by SMAD3-mediated TGF-β signalling are relatively abundant in the SMAD3/SMAD4 binding sequence termed CAGA box (Dennler *et al*, 1998). In the light of ubiquitous expression of SMAD3 in the steady-state condition, future studies are required to identify the factors networking with SMAD3 to specifically induce pDC differentiation.

TGF-β exerts the bidirectional effects on proliferation and differentiation vs. quiescence depending on the HSC subtypes (Blank *et al*, 2015). Extracellular matrix stores and activates latent TGF-β in BM to activate SMAD-mediated TGF-β signalling in HSCs and various hematopoietic progenitor cell populations (Massague, *et al*, 2012; Robertson, *et al*, 2016; Soderberg *et al*, 2009). TGF-β induces HSC hibernation (Yamazaki *et al*, 2009). TGF-β-SMAD3 signalling has been implicated to connect FOXO–p57Kip2 signalling to induce HSC quiescence and self-renewal (Naka *et al*, 2017). Considering the crucial role of SMAD3 in maintaining stem cell quiescence reported in these previous studies, nuclear localization of R-SMADs in freshly isolated BM progenitor cells suggests that the early DC progenitors upstream of MDPs are maintained by SMAD-mediated canonical TGF-β pathway.

In summary, we demonstrate the previously unknown fate determinant roles of SMAD3 as a negative regulator of cDC differentiation and a positive regulator of pDC differentiation in the steady-state. Repression of SMAD3 by phosphorylated STAT3 and c-SKI is required for commitment to cDC, whereas maintenance of SMAD3 via canonical TGF-β signalling is required for pDC differentiation. Although we have narrowed down CD115^+^ CDP and pre-pDC in which SMAD3 expression is regulated for differentiation of cDC and pDC, respectively, our findings would have potential for future studies to identify the unknown lineage-specific DC progenitor cells targeted by SMAD3.

## ACKNOWLEDGEMENTS

We thank Dr. Michael Sporn (Dartmouth Medical College, USA) and Dr. Lalage Wakefield (National Institutes of Health, USA) for helpful discussions and critical reading of the manuscript. We thank Dr. Chuxia Deng (National Institutes of Health, USA) for *Smad3*^*ex8/ex8*^ mice. The project was funded by NRF-2015R1A1A3A04001051, NRF-2018R1A2B6009255 and JSPS KAKENHI JP16K09908 to M.M., JSPS KAKENHI JP17K15735 and NRF-2016R1D1A1B03931785 to JH.Y., JSPS KAKENHI JP18K15252, WISET-2017-083, WISET-2018-737 and NRF-2018R1A6A3A01011885 to E.B., NRF SRC 2017R1A5A1014560 to SH.P. and KHIDI-HI16C1501 to IK.L.

## AUTHOR CONTRIBUTIONS

JH.Y. and M.M. conceived the project idea, designed and performed experiments, analyzed data, and wrote the manuscript. E.B., K.S., M.T., IK.L., JS.H., S.N., T.Y., and T.S. contributed to the animal experiments and drafting the manuscript. Ma.K. and Mi.K. evaluated cytochemical analysis and contributed to drafting the manuscript. JH.J. contributed to the experiments and analyses using flowcytometry and confocal microscopy. SH.P., K.M. and E.B. contributed to the experiments using various mutant constructs and promoter analyses.

## METHODS

Detailed materials and methods are described in Supplemental Experimental Procedures.

